# Hierarchical differences in the encoding of sound and choice in the subcortical auditory system

**DOI:** 10.1101/2022.06.15.496306

**Authors:** Chase A. Mackey, Margit Dylla, Peter Bohlen, Jason Grigsby, Andrew Hrnicek, Jackson Mayfield, Ramnarayan Ramachandran

## Abstract

Detection of sounds is a fundamental function of the auditory system. While studies of auditory cortex have gained substantial insight into detection performance using behaving animals, previous subcortical studies have mostly taken place under anesthesia, in passively listening animals, or have not measured performance at threshold. These limitations preclude direct comparisons between neuronal responses and behavior. To address this, we simultaneously measured auditory detection performance and single-unit activity in the inferior colliculus (IC) and cochlear nucleus (CN) in macaques. The spontaneous activity and response variability of CN neurons were higher than those observed for IC neurons. Signal detection theoretic methods revealed that the magnitude of responses of IC neurons provided more reliable estimates of psychometric threshold and slope compared to the responses of single CN neurons. However, pooling small populations of CN neurons provided reliable estimates of psychometric threshold and slope, suggesting sufficient information in CN population activity. Trial-by-trial correlations between spike count and behavioral response emerged 50-75 ms after sound onset for most IC neurons, but for few neurons in the CN. These results highlight hierarchical differences between neurometric-psychometric correlations in CN and IC, and have important implications for how subcortical information could be decoded.

**New & Noteworthy:** The cerebral cortex is widely recognized to play a role in sensory processing and decision-making. Accounts of the neural basis of auditory perception and its dysfunction are based on this idea. However, significantly less attention has been paid to midbrain and brainstem structures in this regard. Here we find that subcortical auditory neurons represent stimulus information sufficient for detection, and predict behavioral choice on a trial-by-trial basis.

## Introduction

A fundamental question in sensory neuroscience is how psychophysical capabilities of organisms are related to neuronal responses. Early studies characterized this relationship by comparing psychophysical and neuronal measures made at different times (e.g., Barlow et al. 1971; Tolhurst et al. 1983). Studies of the auditory system took this approach, characterizing neuronal responses to a variety of stimuli (Kiang et al. 1965; Goldberg and Brown 1969; Young and Sachs 1979), leading to models of auditory nerve fiber (ANF) response pooling to account for behaviors such as detection (e.g., Siebert 1965; Young and Barta 1985). However, subsequent neurometric analysis, which provides a more direct comparison between neuronal response and behavioral accuracy, demonstrated that thresholds of individual ANFs are much higher than behavioral thresholds (Delgutte 1995), motivating studies of signal detection in the central nervous system. Two candidate structures, the cochlear nuclei (CN) and inferior colliculi (IC), are near-obligate synapses where parallel neural pathways converge (see Oliver and Huerta 1992, Cant 1992, Moore et al. (2010), and Schreiner and Winer (2005) for reviews). Some studies of the IC have identified potential neural codes for detection (e.g., Wang et al. 2021), but these methods (when neural and behavioral measures are collected separately) preclude direct neuronal-behavioral comparison (Parker and Newsome 1998). Almost all previous CN and IC studies exhibit similar limitations, or have not measured performance near threshold (e.g., Ramachandran et al. 1999; Rhode et al. 2010; Ryan et al. 1984; Shaheen et al. 2021; Wang et al; 2021), leaving open fundamental questions about the role of subcortical structures in detection performance.

In contrast, many studies measuring performance at threshold in behaving animals have gained substantial insight into the neural basis of perception (e.g., Britten et al. 1992, 1996; Hernandez et al. 2000; Romo et al. 2002; Purushothaman and Bradley 2005; Nienborg and Cumming 2009; Vazquez et al. 2012). These studies gained insight into two key aspects of responses of sensory neurons: potential neural codes used for task performance, and choice-related activity. Similar studies have been conducted in the auditory system, almost exclusively in cortex, reporting potential neural codes, and correlates of behavioral choice (Christison-Lagay et al. 2017; Francis et al. 2018; Lemus et al. 2009a,b; Niwa et al. 2012a,b), but it is unclear to what degree this information is inherited from subcortical structures due to methodological differences between cortical and subcortical studies. Similar subcortical studies have used a classical conditioning paradigm (Kettner and Thompson 1980, 1985), which involves different circuitry, and provides lower measures of accuracy than operant conditioning tasks (Behrens et al. 2016), or have only measured responses in the CN (Lonsbury-Martin et al. 1987). Thus, it is unclear whether CN and IC represent information sufficient for detection performance, choice-related activity, and if they exhibit hierarchical differences in the encoding of these two variables. This study capitalized on this opportunity by measuring single unit responses in the CN and IC of rhesus macaques performing a reaction time sound detection task at threshold, with the hypothesis that the encoding of sound and choice would improve hierarchically from CN to IC. The findings reveal a hierarchy of sensory and choice-related information in the subcortical auditory system.

## Methods

Experiments were conducted on four male rhesus monkeys (*Macaca mulatta*) that were five (monkeys C, D) and eight (monkeys A, B) years of age at the start of these experiments and were prepared for chronic experiments using standard techniques. All procedures were approved by the Institutional Animal Care and Use Committees at Wake Forest University and Vanderbilt University, and were in strict compliance with the guidelines for animal research established by the National Institutes of Health.

Three surgical procedures were performed to prepare monkeys for this study. A head holder was implanted in the first surgical procedure, and in the next two, craniotomies were performed to access the CN and the IC, and recording chambers were implanted. All surgical procedures were performed using sterile procedure and isoflurane anesthesia. During the first surgical procedure, a head holder (Crist Instruments, Hagerstown, MD) was secured to the skull using bone cement (Zimmer Inc., Warsaw, IN) and 8 mm long stainless steel screws (Synthes Vet, West Chester, PA). The behavioral training on the detection task began after this surgery (see below for details of the task). The other two surgeries implanted recording chambers (Crist Instruments, Hagerstown, MD) on the skull around craniotomies at stereotaxically guided locations. The recording chambers were angled to fit on the skull – the midbrain chamber was tilted lateral 20°, and the brainstem chamber was angled posterior 26° (similar to the approach for the cerebellar flocculus, after Lisberger et al. 1994). The chambers were chosen with bases that fit the cranial curvature, and secured to the skull using bone cement and screws. Pre- and postsurgical analgesics, and, if necessary, antibiotics, were administered, and the monkey was monitored carefully under veterinary supervision until complete recovery had occurred.

All experiments were conducted in double walled sound booths (IAC 1200, Industrial Acoustics Corp., NY, and ER-247, Acoustic Systems, model ER 247) that measured 1.8m x 1.8m x 2m. During experiments, monkeys were seated in an acrylic primate chair custom designed for their comfort and with no obstruction to sounds on either side of the head (Audio chair, Crist Instrument Co., Hagerstown, MD). The monkeys’ heads were positioned 90.1cm in front of speakers placed level with the ears. The speakers (SA1 speaker, Madisound, WI, and Rhyme Acoustics) could deliver frequencies between 50 Hz and 40 kHz, and were driven by linear amplifiers (SLA2, Applied Research Technologies, Rochester, NY) such that the sound level varied less than ±3 dB over the entire frequency range. All calibrations were performed with a 1/4” probe microphone (PS9200, ACO pacific, Belmont, CA) placed at the location of one of the ears of the head-fixed monkey.

### Behavioral task

Stimulus delivery, data acquisition, and reward delivery were controlled by a computer running OpenEx software (System 3, TDT Inc., Alachua, FL). The behavioral task and stimulus delivery mechanisms have been described in detail elsewhere (Dylla et al. 2013; Bohlen et al. 2014). Briefly, the monkeys were trained to detect 200 ms tone bursts with 10 ms rise and fall times. All training involved positive reinforcement. Monkeys initiated trials by pressing down on a lever (Model 829 Single Axis Hall Effect Joystick, P3America, San Diego, CA). When tones were presented after a variable hold time (600 – 1400 ms), monkeys were required to release the lever within a response window, usually extending up to 600 ms after the offset of the tone, for a fluid reward. The response window began with the onset of the stimulus; monkeys were free to respond even before stimulus offset. There were no penalties for not releasing the lever (miss), taken to indicate non-detection. On catch trials (∼20% of the trials), monkeys were required to hold the lever pressed through the response duration, and were not rewarded for a correct reject response. Incorrect releases on catch trials (false alarms) were penalized with a timeout (6 – 10s) in which no tone was presented.

Signals were generated with a sampling rate of 97.6 kHz, and lever state was sampled at 24.4 kHz. Pure tones with sine onset phase, frequency *f*, and amplitude *A* were used as the signal to be detected. The value of *A* was varied such that the sound pressure level of each tone could take values over a 90 dB range, going from -16 dB to 74 dB SPL in 3, 5 or 10 dB steps, and were randomly interleaved. Each tone sound pressure level was repeated 15 to 30 times.

### Neuronal recording methods

A glass coated tungsten electrode (Alpha Omega Engineering, Alpharetta, GA; tip length ∼7-10 µm, diameter ∼5 µm; Thomas Recording GmbH, tip impedance 3 – 4 MΩ) was placed in a guide tube near the surface of the brain. The guide tube was advanced manually ∼10 mm into the brain; the electrode was advanced further into the brain by means of a remotely controlled hydraulic micromanipulator (MO-97, Narishige Inc., Hampstead, NY). The electrode traveled through the cortex to reach the IC, and through the cortex and cerebellum to reach the CN.

As the electrode was driven into the brain, bursts of noise were used as probe stimuli to assess proximity to auditory structures. Proximity to CN or IC was indicated by changes in background responses to the noise bursts. The IC was identified as the auditory structure posterior to the superior colliculus (identified by superficial visual drive, deep layer eye movement sensitivity, and response to visual and auditory stimuli) and anterior to the cerebellum (identified by simple and complex spikes). Further criteria to identify IC (after Nelson et al. 2009) included: (i) short latencies (≤ 20 ms); (ii) reliable, non-habituating responses; and (iii) identification of presence in tonotopic gradient (e.g. Merzenich and Reid 1974). The CN was identified as the region of auditory responses medial to the flocculus (identified by simple and complex spikes, and eye movement sensitivity to ipsiversive and downward eye movements observed over the video monitor).

Once the electrode moved into the CN or IC, single units were isolated using tones (CN: Spirou and Young 1991; Nelken and Young 1994; IC: Ramachandran et al. 1999; Davis 2002). The electrode was advanced through the CN or IC until the signal from the electrode was predominantly from one unit. Single units were verified based on visual inspection and the criterion that >99% of interspike intervals were at least 720 µs (after Nelson et al. 2009).

Once a single unit was isolated, its characteristic frequency (CF: the frequency with the lowest threshold) was manually estimated and used to derive the frequency tuning of the unit via a frequency response map (FRM). A FRM was obtained by measuring the responses of the unit to tones as a function of frequency and sound pressure level. Frequency was varied over a 2 or 4 octave range around the estimated CF in 100 logarithmically spaced steps and at multiple sound levels, starting near CF threshold values, and proceeding in 10 or 20 dB steps, similar to Rocchi and Ramachandran (2020). The FRM was used to estimate the CF of the unit online, which was then used as the frequency at which the monkey performed the detection task. FRMs were also used to classify the response type of the CN or IC units (CN: Evans and Nelson 1973; IC: Ramachandran et al. 1999). Sound pressure levels used for the FRMs ranged from 10 - 15 dB below estimated threshold to as high as 74 dB SPL. The raw waveform of the electrode signal and the waveforms of spikes that exceeded a threshold were sampled at 24.4 kHz and stored for offline analysis.

### Behavioral Data Analysis

Behavioral analyses were based on signal detection theoretic methods (Green and Swets 1966; Macmillan and Creelman 2005), implemented using MATLAB (MATLAB, RRID:nlx_153890; Mathworks, Natick, MA), and have been described previously (Dylla et al. 2013; Bohlen et al. 2014). Briefly, the false alarm rate (*F*) and the hit rate (*H*) for each tone level were calculated based on the number of trials at that sound level with lever releases within the response window and the total number of trials at that sound level. The behavioral accuracy was determined by:

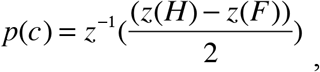

where the *z(H)* and *z(F)* represent hit rate and false alarm rate in units of standard deviation of a standard normal distribution (*z*-score, norminv in MATLAB). The inverse function (*z*^*-1*^) then converts a unique number of standard deviations of a standard normal distribution into a probability correct, a measure of behavioral accuracy (*p(c)*, normcdf in MATLAB). The *p(c)* was calculated for each tone level to create a psychometric function. Since the monkeys’ false alarm rates influenced both the maximum performance and the estimate of chance performance at sound levels below threshold, psychometric functions did not always saturate at 0.5 and 1. To account for that, a Weibull cumulative distribution function (cdf) was fit to the psychometric function:

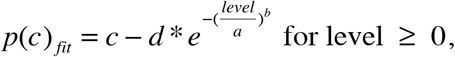

after the methods of Britten et al. (1992) and Palmer et al. (2007), where *level* represents the tone sound pressure level in dB SPL, *a* and *b* represent the threshold and slope parameters, *c* and *d* represent the ceiling and floor saturation rates that could differ from 1 and 0.5, respectively, and *e* is the base of natural logarithms (*e*=2.71828). To account for the sound pressure levels below 0 dB SPL, the data were translated upward by the lowest sound pressure level used, fit with the Weibull cdf, and then the thresholds were translated back to the original sound pressure levels. Threshold was calculated from the fit as that tone level that evoked p(c) = 0.76, and was close to the value of *a*. In all cases, reaction time (RT) was also computed, based on the time of the lever release. RT was computed as follows:

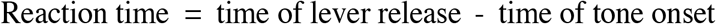

### Neurophysiological Data Analysis

Offline, the neurophysiological data were spike sorted using a Bayesian spike-sorting algorithm (OpenSorter, TDT, Alachua, FL), which were then used for all further analyses. The spike-sorter identified spikes based on a manually set threshold, and then clustered the voltage waveforms exceeding the threshold based on Bayesian classification schemes. The validity of the clustering was further checked based on spike shape and distribution of interspike intervals.

Spikes occurring across the entire 200 ms tone duration in the FRMs were used to classify units into response types (IC: Ramachandran et al. 1999; CN: Evans and Nelson 1973), and to calculate the CF. The calculated CF values matched the online CF estimates to within a sixteenth of an octave more than 98% of the time and did not represent a source of error.

Initial analyses determined the magnitude of the response evoked by each stimulus. For each trial, the response was computed by cumulating the number of spikes evoked by the tone. Thus, a distribution of responses was created for each tone level, including the no tone condition on catch trials. These distributions were used to determine the behavioral accuracy predicted by the ideal observer based on neuronal responses using traditional Receiver Operating Characteristic (ROC) analyses (after Barlow et al. 1971; Tolhurst et al. 1983; Britten et al. 1992; Palmer et al. 2007). Briefly, the distribution of the responses on catch trials were compared to the distributions of responses on signal trials to determine the probability of an ideal observer determining that the responses under the two conditions were different. Such neurometric probabilities were computed at each sound pressure level to generate a neurometric function (after Britten et al. 1992; Palmer et al. 2007). The neurometric functions were fit with Weibull cdf using the equation described above, and neurometric thresholds and slopes were derived from the parameters of the Weibull cdf.

The spiking response on each trial was regressed against the RT on that trial to determine the correlation between response magnitude and RT. Since response latencies can also be a contributor to RT, response latencies were calculated on a trial-by-trial basis using the method described by Rowland et al. (2007). Briefly, spike counts were cumulated from the beginning of the trial, and inflections in the spike count were used to determine response latency and time of response offset on each trial. Both variables were regressed against RT to determine the correlation between a measure of response timing and RT.

The dependence of the neuronal responses to behavioral choice was determined using the choice probability metric (Britten et al. 1996; Cook and Maunsell 2002), and was labeled the detect probability (DP), similar to Cook and Maunsell (2002). When three or more correct and incorrect responses were present, the responses on correct and incorrect trials at the same sound pressure level were compared using ROC analysis. The DP measure calculated the probability with which an ideal observer could predict detection accuracy at that tone sound pressure level based on the trial-by-trial response fluctuations. A DP of 0.5 indicates that the neuronal responses on correct and incorrect trials were not significantly different to an ideal observer, DP > 0.5 indicates that responses were usually larger on correct trials relative to incorrect trials, and DP < 0.5 suggests that responses on incorrect trials were larger than responses on correct trials. Kang and Maunsell (2012) identified potential confounds in calculation of DP (e.g. sample size asymmetry between correct and incorrect responses). Mirroring the methods they described, we sampled neuronal responses 1,000 times with replacement to determine the DP and its significance by permutation test (1,000 repetitions).

When three or more correct and incorrect responses were available at more than one sound pressure level (i.e., the dynamic range was captured by more than one sound pressure level), a grand DP was computed by across all the levels after z-scoring. We also verified that where only analysis of responses at one sound pressure level was possible, the z-scores and response magnitude based ROC analyses were not different from each other.

### Pooling analysis

A pooling model was created to assess the effect of converging CN responses on neurometric performance, to formalize assumptions about how inputs to the IC could lead to responses observed in the IC. The inputs to the model were responses of actual CN neurons recorded. Because the population of units had CF that varied over the audiometric range, unit responses as a function of sound level were normalized such that neurometric threshold reflected the difference between the original neurometric threshold and psychometric threshold. The number of inputs to the model was varied to test the effect of pool size, and determine the pool size that was sufficient to match psychometric performance. The inputs to the model were sampled 1000 times with replacement, and summed, (similar to Britten et al. 1992). The summed responses were subject to ROC analysis described above, to attain neurometric accuracy of the population.

### Statistical analysis of similarity between neurometric and psychometric parameters

Since a goal of this study was to determine neurometric-psychometric correspondences, we used bootstrap methods. The distributions of neuronal and behavioral responses at each tone level were resampled 1000 times to create 1000 distributions of neuronal and behavioral responses at that tone level. This was repeated for each tone level. Thus, the behavioral and neuronal responses as a function of tone level were resampled with replacement to create 1000 different response vs. level functions and hit rate vs. tone level functions. Signal detection theoretic and ROC analyses were performed on each of the resampled hit rate vs. tone level functions and response vs. level functions to derive one thousand neurometric and psychometric functions. Each of the neurometric and psychometric functions was then fit with Weibull cdfs to derive distributions of threshold and slope for psychometric and neurometric conditions. The 95% confidence limits of these distributions (confidence intervals, CI) were then calculated. The CIs were compared across similar psychometric and neurometric parameters for overlap. These distributions were then compared statistically to determine differences in psychometric and neurometric parameter pairs at the 5% level using a Wilcoxon signed rank sum test.

### Histological analyses

All four of the monkeys were euthanized, but we were unable to place markers (lesions or tracer injections at physiologically characterized sites). The brains were extracted (fresh in the case of monkey A and post fixed, and after transcardial perfusion with 4% paraformaldehyde following euthanasia and exsanguination in other monkeys). The subcortical regions were blocked caudal to the medial geniculate nucleus to include both the IC and CN. The brain blocks were allowed to sink in a 30% solution of sucrose in phosphate buffer (pH 7.2 – 7.4) and cut into 50 µm sections. The sections were stained with cytochrome oxidase (CO, Millipore Inc., Billerica, MA) and then mounted on slides, or mounted on slides and stained with thionin (Sigma, St. Louis, MO). The thionin stained sections were used to identify subdivisions of the CN and the IC. Figure 1 shows sections through the IC and CN in monkeys A and B stained with CO (an example from monkey D is shown in Rocchi and Ramachandran (2020)). Tracks were found in both structures (ovals in Figure 1) and confirmed based on adjacent sections. Fewer tracks were found in the CN than in the IC, but could be attributed to the 26° anterior-posterior electrode angle being viewed through nearly coronal sections. Tracks were found throughout the rostrocaudal extent of the IC and the CN, including tracks through the dorsal CN (DCN), providing anatomical confirmation that the recordings in these macaques were indeed from the IC and the CN. The tracks in the IC were found passing through the central nucleus, consistent with the recording approach of looking for tuned auditory responses. Tracks were found to extend throughout the dorsoventral extent of the central nucleus of the IC, indicating that the electrodes traversed a majority of the representation of the frequency extent, consistent with the range of unit CFs measured (see Figures 4A and 4B). Electrode tracks were also found throughout the anterior-posterior extent of the CN, indicating evidence of sampling throughout the frequency extent in the CN also (also see Figures 7A and 7B).

**Figure 1.**
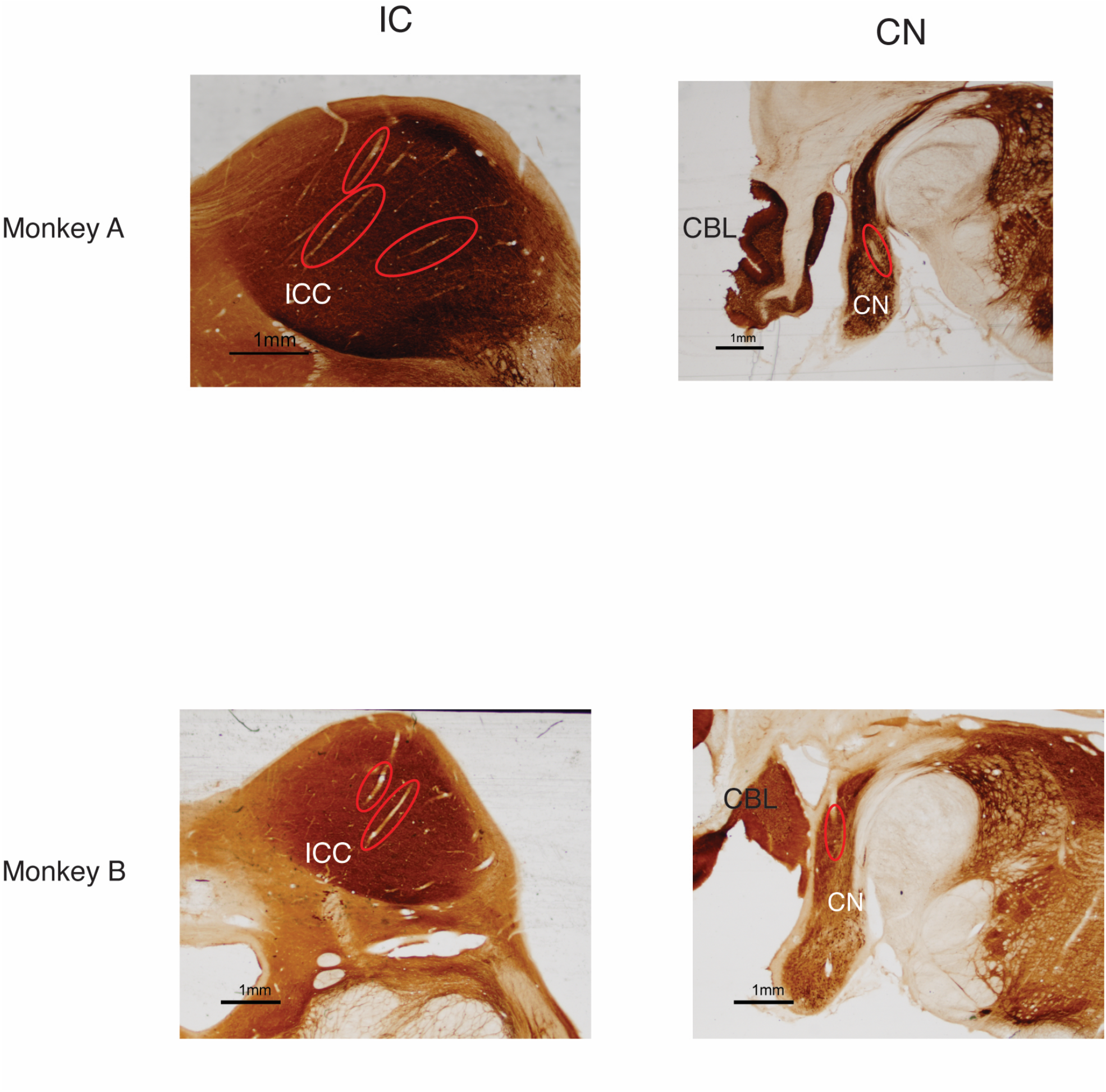
Histological sections from two monkeys. Examples of coronal sections through the CN and IC. Sections are shown from the brains of two monkeys (Top: monkey A, bottom: monkey B). The columns show sections from the IC (left) and the CN (right). Red ovals outline the electrode tracks at that location. Scale bar (1mm) is shown in each panel.

## Results

### Basic response properties

The responses of 108 single units with complete psychometric and neurometric data were studied in the IC of four monkeys (A, n=25; B, n=36; C, n=9; D, n=38). Responses were also obtained from 92 single units with complete psychometric and neurometric functions and sufficient number of trials in the CN in the same monkeys (A, n=29; B, n=44; C, n=1; D, n=18). As observed in multiple species, the IC had a tonotopic region in which the CF of single units increased progressively as the electrode traveled dorsolateral to ventromedial (Reviewed in Oliver and Huerta (1992); Schreiner and Winer (2005); and Moore et al. (2010)); the CN had multiple tonotopic fields. All the IC data presented in this report were obtained in the tonotopic region of the IC, most likely the central nucleus of the IC; these were confirmed by the majority of tracks traversing through the central nucleus of the IC (see Figure 1). The physiological signatures (short latency auditory responses, the tonotopic gradient, and the FRM types) were consistent with the recordings being from the central nucleus of the IC (ICC), and from all divisions of the CN. The majority of the CN units were obtained in the anterior two-thirds of the structure (putative ventral CN).

The CFs of the units used in this manuscript ranged between 200 Hz and 32 kHz for IC units and between 300 Hz and 34 kHz for CN units; the sample of IC units in monkeys C and D had predominantly low CFs because those monkeys were also used in studies of neuronal encoding of vowels, which are lower frequency signals. IC units were classified, based on their binaural diotic frequency response map (FRM) as IC type V (V-shaped excitatory FRM about CF, n=68) or IC type I (V-shaped excitatory FRM about CF, with inhibition flanking excitation, n=35) units (after Ramachandran et al. 1999). The responses of a small number of type O units (n=5, after Ramachandran et al. 1999), whose responses to CF tones at high sound pressure levels fell below spontaneous activity, were recorded, consistent with previous reports (Nelson et al. 2009). CN units were classified into CN type I units (the Roman numeral 1, having V-shaped excitatory FRMs with no inhibitiory responses, n=44), CN type III units (Roman numeral 3, V-shaped excitatory FRM and flanking frequency regions of inhibition, n=37), CN type IV units (Roman numeral 4, low sound level excitation at CF, and inhibitory responses at higher levels, n=6), and CN type V units (Roman numeral 5, inhibition at all responsive frequencies and sound levels, n=4) based on existing classification methods (e.g., Evans and Nelson 1973; Shofner and Young 1985).

Figure 2 shows some exemplar waveforms and some basic response measures in the IC and the CN. Figure 2A shows waveforms obtained during recordings of a single unit in the IC and one during the recordings of a single unit in the CN. As expected for single units, the amplitude was very stable over time for all units classified as single units. Figure 2B shows the distribution of baseline/spontaneous activity (activity when sound stimuli were not presented) in the IC and in the CN (filled and open bars respectively). The mean spike count over the 200 ms stimulus duration on catch trials were calculated and scaled to provide the spontaneous firing rates. There was not a significant difference in the spontaneous rates depending on unit type for either the CN or the IC (Kruskal Wallis test, *p*>0.05), so the spontaneous rates were combined across unit types. The spontaneous rates of CN and IC units took values over the similar ranges (IC: 1.8 – 60.2 spikes/s; CN: 1.2 – 69.8 spikes/s), but their distributions were very different.

**Figure 2.**
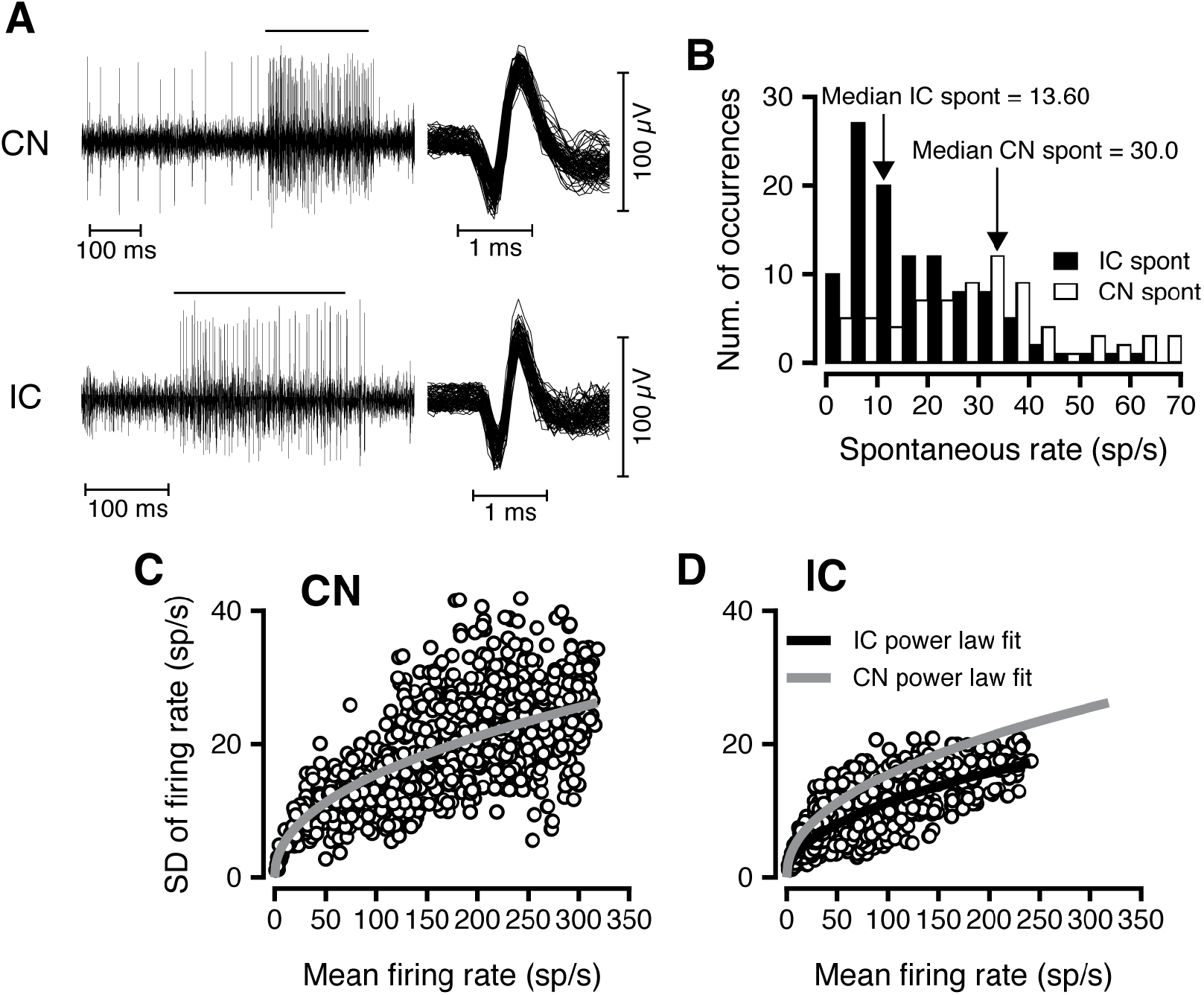
Response statistics in the CN and IC. A. Examples of spike trains and spike snippets. Left: Spike train in one single trial around CF tone presentation is shown for an exemplar IC unit (top), and CN unit (bottom). The duration of stimulus presentation is shown by the horizontal line, and results in greater frequency of spike occurrence in both spike trains. Right: Spike snippets from the IC and CN spike trains. B. Distribution of firing rate in the absence of tones (spontaneous rates) in the CN (open bars) and IC (filled bars). The spontaneous rates were calculated from the spike counts on catch trials (see Methods). C. Relationship between the standard deviation (SD) and mean of the firing rates for CN units. The thick grey line represents the best power law fit to the data (y=1.85*x^0.47^). D. Relationship between mean and standard deviation of firing rates for IC units. The black line represents the best power law fit to the IC data (y=0.85*x^0.46^). The grey line is the power law fit for CN units, shown for ease of comparison.

Spontaneous rates of IC units tended to be lower (mean = 18.6 spikes/s, median =13.65 spikes/s). CN unit spontaneous rates tended to be higher, with a mean of 30.3 spikes/s, and a median of 30 spikes/s. The ranges of spontaneous firing rates of CN and IC units were consistent with the published values or the ranges in unanesthetized animals (e.g., IC: Ramachandran et al. 1999; CN: Evans and Nelson 1973). The spontaneous activity in the CN and IC were significantly different from each other (Wilcoxon ranksum test, Z=5.53, *p*=1.8×10^−8^).

The variability of neuronal responses is a crucial determinant of stimulus encoding. Figures 2C and D show the relationship between the standard deviation (SD) of the firing rate and the mean firing rate in response to the tones at CF for CN and IC units, respectively. Both figures show that variability of the firing rate was related to the mean, consistent with findings in the auditory system (e.g., May and Huang 1997; Reiss et al. 2011). For both CN and IC, the relationship between mean and standard deviation was best fit with a power law function (grey and black curves in Figures 2B and C respectively). However, there are differences between the CN and IC. The maximum mean response of IC units (maximum = 260 spikes/s) was lower than that of CN units (maximum = 314 spikes/s). More noteworthy was that the standard deviation (or variability) of IC firing rates was lower than that of CN firing rates. Thus, CN units had higher baseline activity, and higher variability in their activity than IC units. Thus, exceeding the baseline based on statistical criterion would take larger changes in the mean firing rates of CN units as compared to IC units, suggesting that neurometric thresholds of CN units would be higher than that of IC units.

### Responses in the inferior colliculus (IC)

Behavioral thresholds to tones were obtained from all four monkeys prior to neurophysiological measurements. The audiometric thresholds of the monkeys have been published in earlier papers (Dylla et al. 2013; Bohlen et al. 2014) and were not different from the published audiometric functions from other labs (e.g., Stebbins et al. 1966; Pfings et al. 1975). The psychometric thresholds during physiological recordings did not differ from the baseline psychometric thresholds obtained prior to chamber implantation and neurophsyiological measurements (Dylla et al. 2013). The responses of IC type O units were excluded from analysis relative to detection behavior due to their posited role in sound localization (Davis et al. 2003).

Figure 3 shows the responses of single units recorded in the IC of awake and behaving monkeys to CF tones as a function of tone level, and the calculation of neurometric functions and parameters. Figures 3A and B show spike count as a function of the CF tone sound pressure level for an IC type V (CF=2.2 kHz) and IC type I unit (CF=25.6 kHz). The relationship between spike count and tone level for these unit types could be monotonic or nonmonotonic, but never fell below spontaneous rate (solid line, Figure 3A and B). The distribution of responses at each tone level was compared with the responses on catch trials (colored and white bars, respectively, Figures 3C and D). At low tone levels (e.g., -6 dB SPL), the units were unresponsive to the CF tone, with responses during stimulus duration matching those on catch trials when no tone was presented. At high tone levels (e.g., 54 dB SPL), the units’ responses to the CF tone exceeded the magnitude of the responses on catch trials. The increasing responsiveness to the tone created separation between the catch trial and the signal response distributions (Figure 3C and D). A receiver-operating characteristic (ROC) was created by systematically defining criteria over the range of spontaneous and signal response distributions to create false alarm and hit rates. The area under the ROC curve is an ideal observer’s probability of being correct in a binary choice task (tone present vs. not present), based on the neural firing rate during the tone (“neurometric”, open symbols, Figure 3E and F). These neurometric functions were fit with Weibull cdfs (dashed curve, Figure 3E and F) as described in the Methods section. To directly compare these with behavioral performance, simultaneously measured psychometric functions (filled circles) and their Weibull cdf fit (solid curve) are also shown in the same panels. The sound pressure level that evoked *p(c)*_*fit*_ = 0.76 was considered the threshold (solid and dashed vertical lines indicate psychometric and neurometric thresholds respectively, Figures 3E and F).

**Figure 3.**
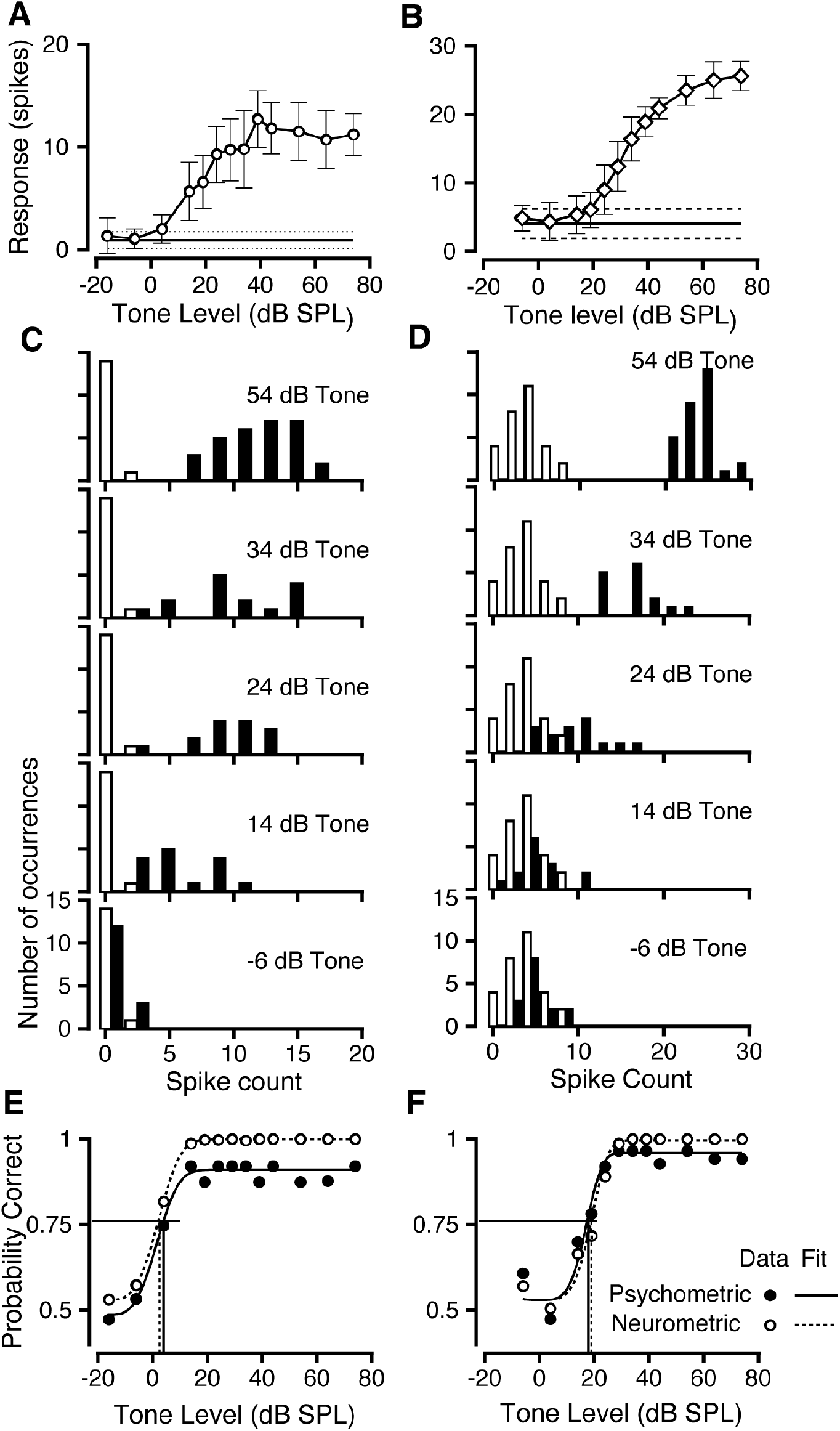
Responses of representative type V (CF=2.2 kHz) and type I units (CF=25.6 kHz) in the IC of the behaving primate to tones at CF as a function of sound pressure level, and ROC analysis. A, B. Responses (spike count) to a tone at CF as a function of sound pressure level for the type V (A), and type I unit (B). Error bars represent standard deviation of the trial-by-trial spike count. Spontaneous activity is shown as a horizontal line (the range, shown by dashed lines, represents ± 1 standard deviation), and was computed from the spike counts evoked on catch trials. C, D. Distributions of evoked spike counts on catch trials (white bars) and signal trials (filled bars) at various sound pressure levels indicated for the exemplar type V (C) and type I unit (D). E, F. Neurometric functions (open circles) based on the response distributions in panels C and D respectively, plotted along with the psychometric function (filled circles) obtained simultaneously. Solid and dashed lines represent Weibull cdfs fit to the psychometric (solid)) and the neurometric functions (dashed). Solid and dashed vertical lines show psychometric and neurometric thresholds, respectively.

The neurometric functions for the exemplars were remarkably similar to the psychometric functions (IC type V: neurometric: *a*=20.77, *b*=3.32; psychometric: *a*=19.77, *b*=3.28; IC type I: neurometric: *a*=24.6, *b*=4.45; psychometric: *a*=26.8, *b*=4.5). In both exemplar cases, the neurometric and psychometric parameters (thresholds and slopes) were not significantly different from each other, determined via Monte-Carlo methods (see Methods; IC type V: threshold: *p*= 0.427; slope: *p*=0.226; IC type I: threshold: *p*= 0.513; slope: *p*= 0.39). The similarity between the psychometric and neurometric parameters was a very common feature of the data obtained in the IC.

Figure 4 plots thresholds against the characteristic frequency for all IC type V and I units across all the monkeys. Figure 4A (left) plots the psychometric thresholds against the frequency of the target tone (the CF of the unit being recorded). The different symbols show the data from the different monkeys. The data shows that units with a large range of CFs were recorded in all monkeys, that the behavioral thresholds showed a U-shaped contour that has been observed in audiograms of macaques (e.g., Stebbins et al. 1966; Dylla et al. 2013). Figure 4A (right) shows neurometric thresholds as a function of unit CF for monkeys A – D using symbols similar to the behavioral panel. The neurometric thresholds also fell along a U-shaped contour similar to the psychometric threshold, though IC neurometric thresholds took a larger range of values. To more easily visualize differences in psychometric and neurometric thresholds, we plotted the difference between psychometric and neurometric threshold for each unit against the unit CF (Figure 4B). The large number of units with low CF (< 1 kHz) reflects data collected during a study of the neuronal representation of vowels (low frequency sounds). The figure shows that the observation of similarity of psychometric and neurometric thresholds were not restricted to a single frequency range but were a general feature across CFs. Figure 4B also revealed that some units had higher thresholds than behavior.

**Figure 4.**
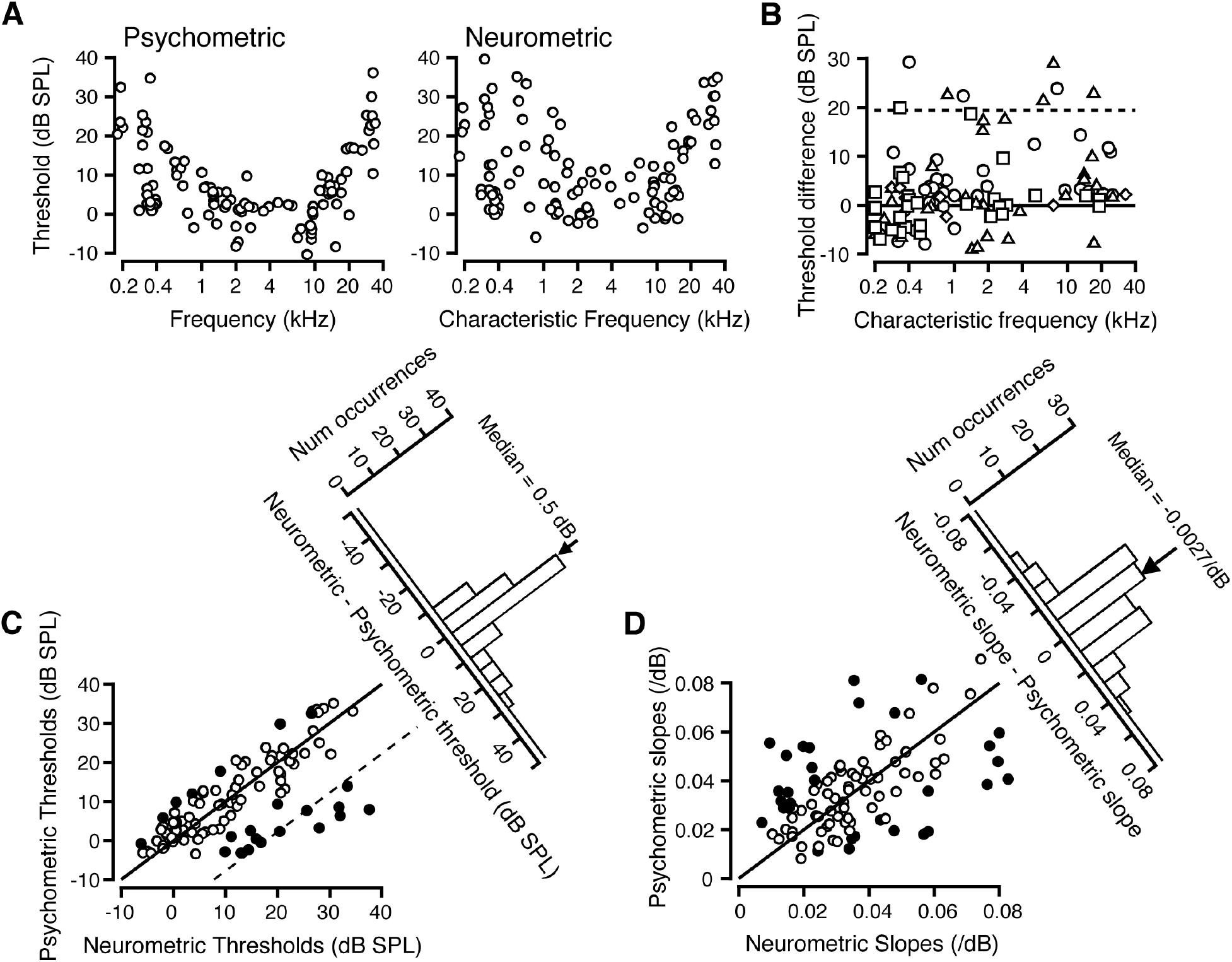
Comparison of psychometric and neurometric parameters in the IC. A. Plot of psychometric thresholds (left) and neurometric thresholds (right) in relation to unit CF. Each circle represents the data obtained while recording the responses of one single unit tuned to the behavioral target tone frequency. B. The difference between psychometric and neurometric threshold is plotted against the CF of the unit. Various symbols represent data from different monkeys. C. Comparison of psychometric and neurometric thresholds. Each symbol represents neurometric threshold from one unit, and the psychometric threshold obtained simultaneously. Filled symbols indicate psychometric and neurometric thresholds significantly different from each other. A histogram of the threshold differences (neurometric – psychometric threshold) is shown perpendicular to the diagonal. D. Comparison of psychometric and neurometric slopes. The format is the same as in Figure 4C. The slope difference histogram is shown perpendicular to the diagonal.

Figure 4C plots IC neurometric thresholds against simultaneously measured psychometric thresholds. There were no significant differences in the relationship between psychometric and neurometic thresholds as a function of unit type, so units of both major types are shown together. Filled and unfilled circles indicate units whose neurometric thresholds were and were not significantly different from the psychometric threshold (see Methods), respectively (Figure 4C). The clustering of circles around the diagonal indicates that neurometric thresholds for many IC neurons were around and not significantly different from psychometric thresholds. There was a significant correlation between individual neurometric and psychometric thresholds (r=0.625, *p*<0.01), strengthening the idea that the strong correlation is at the single unit level, not just on the average. Such correlations could arise out of common factors. Psychometric thresholds in primates vary with tone frequency (e.g., Green et al. 1966; Pfingst et al. 1975; Dylla et al. 2013), as do neuronal thresholds in various auditory regions (e.g., IC: Ramachandran et al. 1999). When multiple regression was used with CF as covariate, the partial correlation between psychometric and neurometric threshold was much reduced and not significant (r=0.155, *p*=0.119). The range of neurometric thresholds observed in Figure 4C is due to the sample containing units with CFs that spanned nearly the entire audiometric range, and thus showing a threshold ranging from ∼-5 to 35 dB SPL. The histogram of threshold differences shows a strong mode around 0, and ∼75% of units (76/102) had threshold differences within ±10 dB (Figure 4C). The median was close to zero (0.5 dB): on the average, the psychometric and IC neurometric thresholds were similar in value, and were not significantly different from each other (Wilcoxon signed rank sum test, Z=0.89, *p*=0.374).

For IC unit responses to adequately represent behavioral performance, IC neurometric functions need to account for both threshold and shape (slope) of the psychometric functions (Parker and Newsome 1998). Since the slope parameter of a Weibull cdf depends on threshold, the slope parameter *b* was not used; rather, the slope was measured as the linear slope of the dynamic portion of the neurometric or psychometric function (10 – 90% of the range). Figure 4D shows a comparison of the psychometric and neurometric function slopes. The data points are scattered around the diagonal line of equal slopes, and with much more variability than that observed for thresholds (compare Figures 4C and D). As with thresholds, the relationship between psychometric and neurometric slopes was independent of CF and response type. As with thresholds, resampling methods were used to compare individual psychometric and neurometric thresholds, and units with significantly different slopes are shown with filled circles. The psychometric and neurometric slopes were significantly correlated (*r* = 0.497, *p* < 0.05), suggesting good match between the two slopes at the single unit level. The slope difference histogram reflects the same trend by a broad distribution centered on 0 (median = -0.002/dB).

### Responses in the cochlear nucleus (CN)

The strong match between the responses of IC units and behavioral responses could be due to either: (i) behaviorally matched neuronal responses that input to the IC; or (ii) synthesis in the IC, *de novo* or via cortical efferents. To separate the two possibilities, responses were obtained in the CN during the same task. Since the responses were uniform across locations within the CN, the data were combined across all CN recording locations. As in the case of the IC, psychometric thresholds during neurophysiological experiments did not deviate significantly from baseline psychometric thresholds at similar frequencies.

Figure 5 shows the responses of an exemplar CN type I unit (Figures 5A, C, E) and CN type III unit (Figures 5B, D, F) and the ROC analysis in a manner similar to Figure 2. The response magnitudes of CN were monotonic or weakly nonmontonic as a function of CF-tone sound pressure level (Figures 5A and B). As with IC units, the distributions of CN responses on signal and catch trials (Figures 5 C, D) were converted, using ROC analysis, into neurometric functions (Figures 5 E and F). The neurometric functions (open circles) were fit with a Weibull cdf (dashed curve), and are shown along with the simultaneously obtained psychometric function (filled circles) and its Weibull cdf fit (solid curve). Threshold and slope parameters were extracted for both data sets. In both examples, the psychometric and neurometric functions appeared very different and the fit parameters were different (CN type I: neurometric: *a*=4.2, *b*=3.43; psychometric: *a*=-2.7, *b*=4.94; CN type III: neurometric: *a*=25.1, *b*=6.17; psychometric: *a*=17.7, *b*=4.8). The psychometric and neurometric thresholds for the exemplar CN units were significantly different from each other as assessed by Monte-Carlo methods (see Methods; CN type I: *p*= 0.0134; CN type III: *p*= 0.009). However, the psychometric slopes for the type I unit were significantly different from (higher than) the neurometric slope (slope: *p*= 0.0099), the psychometric and CN type III neurometric slopes were not significantly different (slope: *p*= 0.319).

**Figure 5.**
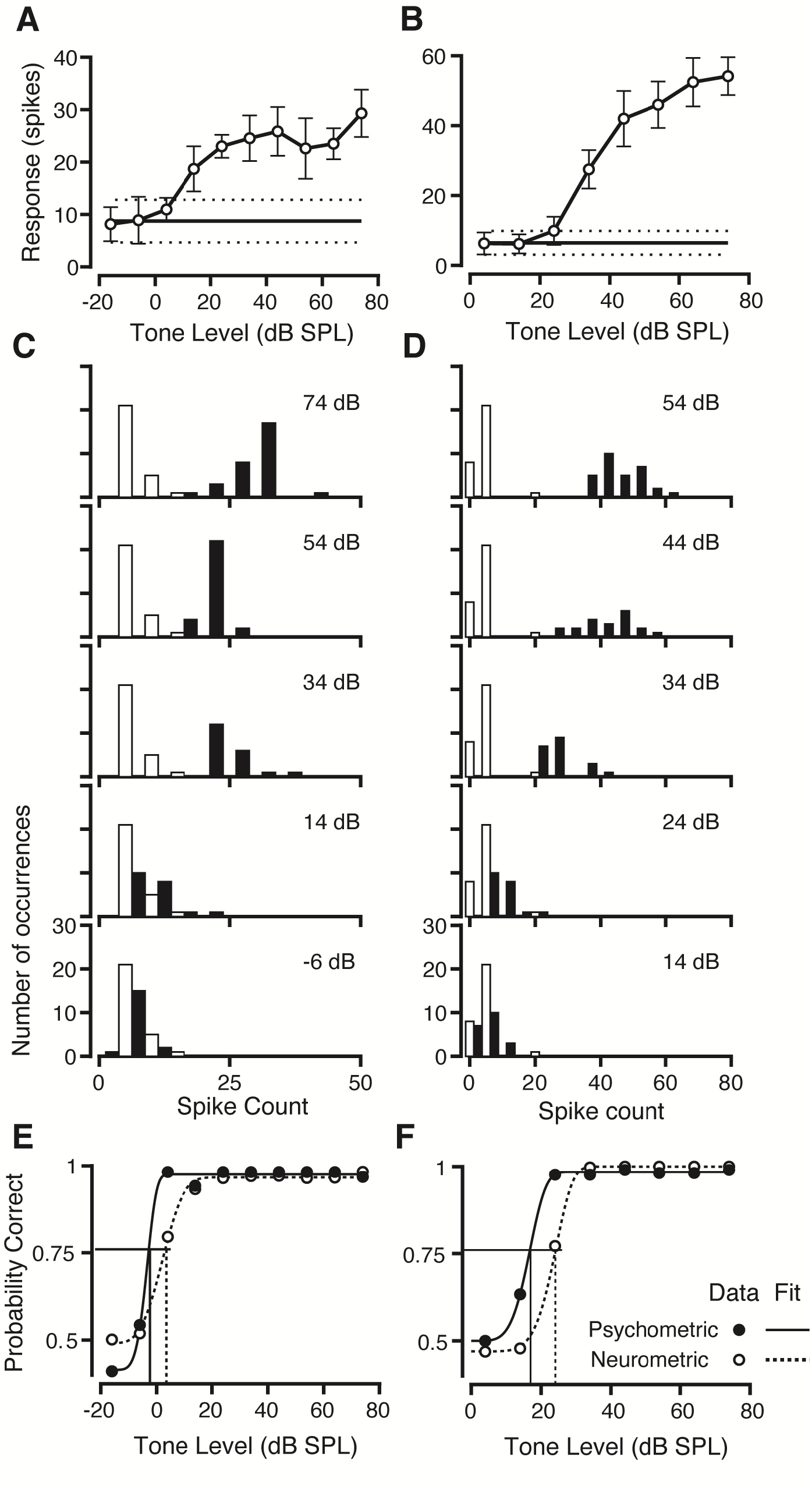
Responses of units in the CN to tones at CF as a function of sound pressure level, and results of ROC analysis for type I (CF=2.4 kHz) and type III (CF = 32 kHz) units. The format is the same as Figure 3. A, B. Responses (spike count) as a function of tone sound pressure level for a CN type I (A), and CN type III unit (B). C, D. Distribution of evoked spike counts on catch trials (unfilled bars) and signal trials (filled bars) at the indicated tone sound pressure levels for the CN type I (C) and the CN type III units (D). E, F. Neurometric functions based on the distribution of responses in panels C and D (open circles), plotted along with the psychometric functions obtained simultaneously (filled circles) for the CN type I unit (E) and CN type III unit (F). Weibull cdf fits for the neurometric (dashed) and psychometric functions (solid) are also shown. Vertical lines show psychometric (solid) and neurometric thresholds (dashed).

Figure 6 shows a comparison of psychometric and neurometric parameters while recordings were made in the CN using a format similar to Figure 4. Figure 6A (left) shows behavioral thresholds during recordings in the CN. The threshold trends were similar to those in Figure 4, and are in general, similar to macaque audiograms (e.g., Stebbins et al. 1966; Pfingst et al. 1978; Dylla et al. 2013). Figure 6A (right) shows CN neurometric thresholds as a function of CF. Thresholds did not vary with the identity of the monkey, and can be seen by comparing the relative locations of the different symbols. In contrast to the IC data in Figure 4A(right), the CN neurometric thresholds were biased to higher tone levels (Figure 6A, right). The differences between psychometric and neurometric threshold were plotted against the unit CF for easy visualization (Figure 6B, same format as Figure 4B). Threshold differences between psychometric and neurometric functions in the CN and IC were significantly different (Wilcoxon ranksum test, Z=8.831, *p*<0.00001). The difference between psychometric and neurometric thresholds for CN units (circles) were not related to the CF of the unit, and across the entire range of frequencies sampled, units in the CN had neurometric thresholds that were higher than psychometric threshold (Figure 6B). Note that the psychometric-neurometric threshold differences for the majority of CN units overlapped with those of the population of IC units with the higher threshold differences (grey shaded region).

**Figure 6.**
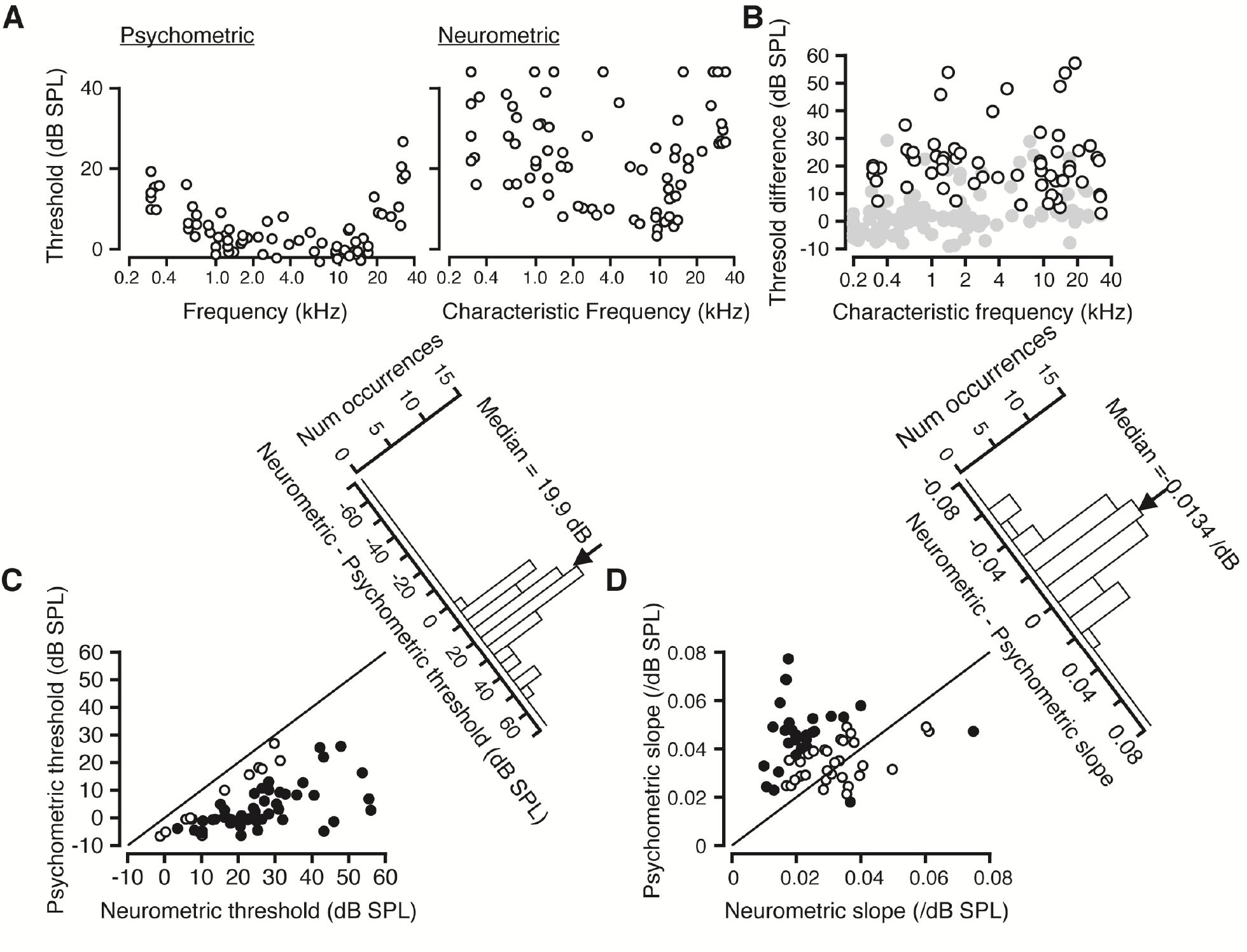
Comparison of psychometric and CN neurometric parameters. Figure uses the same format as Figure 4 and Figure 5B. A. Left, psychometric thresholds plotted against tone frequency. Right, CN neurometric thresholds calculated from responses obtained simultaneously with behavior. Some units have neurometric thresholds higher than 40 dB SPL, and these are shown in a line above the ordinate axis. B. Threshold differences (psychometric – neurometric) vs. the CF of CN units. Format is the same as Figure 4B. IC data (Figure 4B) is shown in grey for comparison. C. Comparison of psychometric and neurometric thresholds of CN single units, and a threshold difference histogram, using the same format as Figure 4A. D. Comparison of psychometric and neurometric slopes, and a slope difference histogram, in the same format as Figure 4C. E. Histogram of DP; format of figure is the same as Figure 5B. Black and grey filled bars represent units with DP values significantly different higher than and lower than 0.5, respectively.

The neurometric-psychometric comparisons were performed for all CN units, and the results shown in Figure 6C and D, similar to Figures 4C and D. Resampling methods were used to determine differences between psychometric and neurometric parameters in a manner similar to that described for IC units. Figure 6C shows a comparison of psychometric and neurometric thresholds, using the same format as Figure 4C. Consistent with the exemplars shown in Figure 6 and panels 6A and B, the entire population fell under the diagonal line of equal thresholds (solid line), and only a small number of CN units had neurometric thresholds that did not differ significantly from psychometric thresholds (*p*>0.05; open circles, Figure 6C). CN neurometric thresholds were, on average, higher than psychometric thresholds (median threshold difference = 19.8 dB), and significantly different from psychometric thresholds (Wilcoxon signed ranksum test, Z=3.41, *p*= 6.5×10^−4^). Despite most threshold pairs being significantly different, the psychometric and neurometric thresholds were still correlated (r=0.46, *p*<0.05), suggesting that even though psychometric and neurometric thresholds were different in magnitude, higher (lower) psychometric thresholds were measured along with higher (lower) CN neurometric thresholds. However, much like the IC data, these were based on the relationship between the thresholds and unit CF; thus the partial correlation between psychometric and neurometric thresholds while using CF as a covariate was not statistically significant. Further, the variance of the CN thresholds (re: psychometric thresholds) were significantly different from the variance of the IC neurometric thresholds (re: psychometric thresholds (Ansari Bradley test, W=4398, W*=2.5685, *p* = 0.0078).

A comparison of the slopes of the psychometric and CN neurometric functions is shown in Figure 6D. In spite of the dichotomy of the exemplars shown in Figure 5, across the population, there was no segregation of type I and type III units slopes and their relationships with the psychometric slope. A larger proportion of CN neurometric slopes were significantly different from the psychometric slope relative to the IC (compare filled circles, Figures 4D and 6D). On the average, CN neurometric slopes were shallower than the psychometric slopes (median difference=-0.014/dB); the difference between the neurometric and psychometric slopes for CN units was significantly different from zero (Wilcoxon signed ranksum test, Z=2.91, *p*=3.61×10^−3^). The differences between psychometric and neurometric slope in the CN and the IC were significantly different from each other (Wilcoxon rank sum test, Z=2.08, *p*=0.075), suggesting that the neurometric sensitivity to tone level differences was larger in the IC than in the CN.

### Detect probabilities (DP)

Early studies posited that for unit responses to be related to perception, the variability in neuronal responses should be related to behavioral variability on a trial-by-trial basis (Britten et al. 1996; Parker and Newsome 1998). More recently, studies have provided evidence that these relationships reflect interneuronal correlation structure (Liu et al. 2013; Zaidel et al. 2017). In this study the relationship between neuronal and behavioral variability was quantified using the detect probability measure (DP, see Methods; after Britten et al. 1996; Cook and Maunsell 2002; Kang and Maunsell 2012), which provides an estimate of the probability that an ideal observer could predict behavioral accuracy based on spiking activity. Figure 7A shows the DPs for all IC units. About 75% of IC units (n=81/108) showed DP values that were significantly different from 0.5, and thus significant response differences related to behavioral outcome; most of these were units that had DP values greater than 0.5 (black bars; responses that were higher on correct trials relative to incorrect trials), and the rest were units that showed DP values less than 0.5.

**Figure 7.**
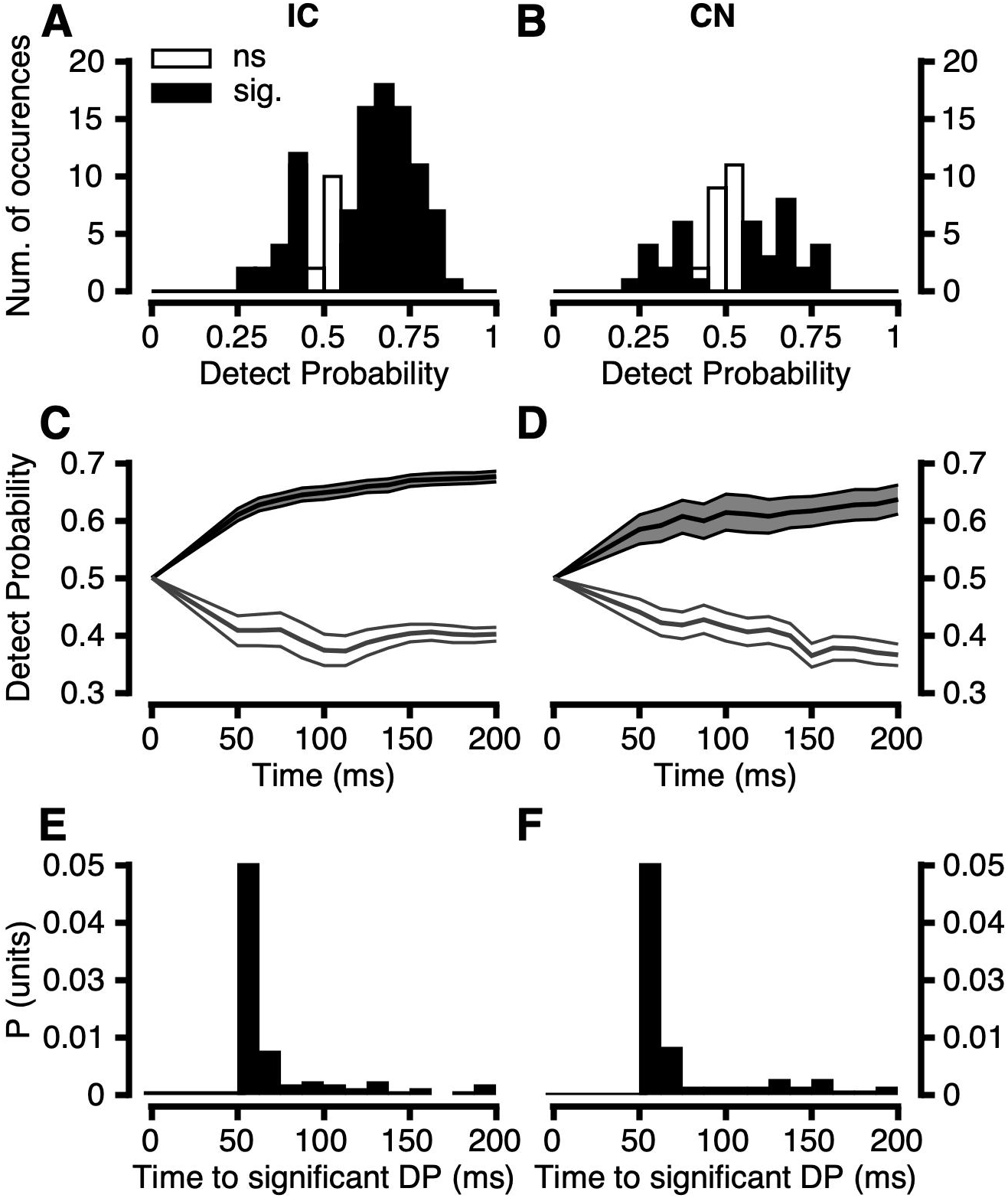
Relationship between fluctuations in spike count and variations in behavioral response. A,B. Histogram of detect probability (DP, after Britten et al. 1996; Cook and Maunsell 2002, see methods for calculation) for all units in the IC (A) and in the CN (B). Black filled bars represent units that showed a DP significantly different than chance (0.5), as assessed by permutation test (see Methods). White filled bars represent units with DP values not significantly different from chance. C,D. Detect Probability (Mean +/- SEM of all units) as a function of time after tone onset for all units in the IC (C) and the CN (D). Some units’ DP increased over time (black error-bands) while others decreased DP over time (white error-bands). E,F. Proportion histograms of time elapsing between tone onset and significance of DP in the IC (E) and CN (F).

Significance of each DP was determined by resampling (see Methods), followed by permutation test, after Kang and Maunsell (2012). The median DP value was 0.647, while the mean was 0.62, suggesting that correct trials were correlated with larger total responses in the IC. DP values were calculated for CN units, and their distribution is shown in Figure 7B. The distribution of DP for CN units was centered closer to 0.5 than the corresponding distribution for IC units. Smaller proportions of units showed DP values significantly different than 0.5: some units (n=23, 25.3%) showed DP values greater than chance (increased responses on correct trials, black filled bars), while others (n=14, 15.3%) showed DP values less than 0.5 (corresponding to decreased responses on correct trials relative to incorrect trials, black bars). The median value of DP for CN single units was 0.53, not significantly different than chance (permutation test, *p* = 0.11). The distributions of DPs in the IC and CN were significantly different, as assessed by a 2-sample Kolmogorov-Smirnov test (p = 0.003).

Many studies have measured CP or DP as a function of time (response duration) to gain insight into the origin of choice related activity in sensory neurons (see Discussion). In this study, DPs were calculated using different time-windows, beginning at tone onset. Figure 7 C-D depicts the mean (+/- SEM) DP across the population in the IC (Figure 7C) and CN (Figure 7D). DPs were concentrated at 0.5 at the beginning of the response, and diverged from chance within the first 75 ms, on average. Some units increased DP over time (black error bands in Figure 7C and D), while others decreased DP over time (white error bands). To more closely inspect this trend, the time required to reach a significant DP was calculated for each neuron exhibiting a significant DP. Most neurons exhibiting a significant DP in the IC (Figure 7E) and CN (Figure 7F) reached significance in the first 100 ms. This is apparent in the proportion histograms, whose peak bins occur 50-100 ms. The mean time to significance was 64.9 ms in the IC, and 67.8 ms in the CN. The distributions of time-to-significance values were not significantly different between brain regions (Kolmogorov-Smirnov test, *p* = 0.9).

### No trial by trial correlation between subcortical neuronal responses and reaction time

Behavioral reaction times (RTs) decreased as the tone sound pressure level increased (e.g., Green et al. 1966; Dylla et al. 2013). Figure 8 shows the relationship between response magnitudes and RT for CN and IC units. Figures 8A and D shows how RT co-varied with the responses for the exemplar IC unit from Figure 2A and the exemplar CN unit in Figure 6A, respectively. In general, RTs decreased as the response magnitude increased. A linear fit best captured the variability of the data (shown by the black line), and its slope was termed the neurometric RT slope. For most units (IC: 96%; CN: 96%), the neurometric RT slopes were significantly different from zero (*p*<0.05, as assessed by Monte-Carlo methods, see Methods), suggesting a relationship between response magnitude and RT. However, when the variations in RT and response magnitude were compared on a trial-by-trial basis at each sound pressure level, the two parameters were not significantly correlated (*p* > 0.05, examples shown in Figure 8B and 8E). Thus the relationship between RT and response magnitude for CN and IC neurons may be an epiphenomenon of the relationship between response magnitude and tone level. Similar results were obtained in analyses of latency. Latencies were longer at lower sound levels (when RTs were longer), and shorter at higher sound levels (when RTs were shorter). A linear fit could account for the relationship between latencies and RTs (similar to figures 8A and C), but there was no significant relationship between RTs and the related neurophysiological latencies on a trial-by-trial basis at each sound level (Figure 8B and 8E). Consistent with these findings, multiple regression between response magnitude (or latency) and reaction time including sound pressure level as a covariate resulted in partial correlations that were not statistically significant (p>0.05). The distribution of the neurometric RT slope (Neurometric RTS) for IC and CN single units are shown in Figures 8C and F. All slope values were negative, indicating that RT decreased as IC and CN responses increased for all units in this population, with a median rate of -3.1 ms/spike. The distribution of neurometric RT slopes for CN units (shown in Figure 8F) was not significantly different than that found for IC units (Wilcoxon ranksum test, *p*>0.2).

**Figure 8.**
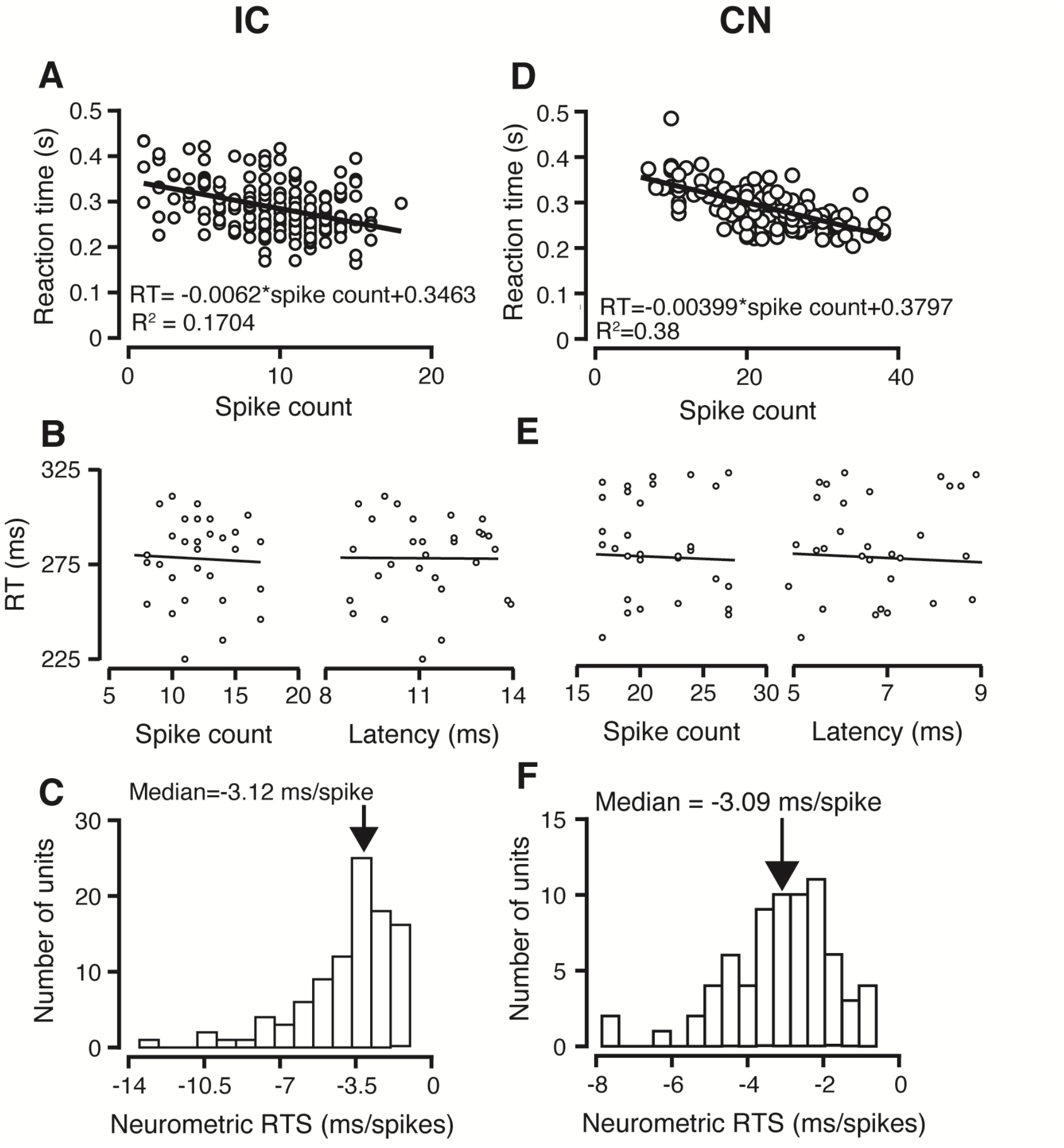
Neuronal correlates of RT in the IC and CN. A, D. Behavioral RT vs. spike count evoked in response to suprathreshold tones for the representative IC type V (A) and CN type I unit (D) whose responses are shown in Figures 2A and 6A. Each symbol represents the RT and the neuronal response on a trial resulting in a lever release. The line represents a linear fit to the data (slope: neurometric RT slope). B, E. Shows RT as a function of spike count (left) and latency (right) at a fixed sound level in the IC (B) and the CN (E). C, F. Histogram of neurometric RT slope for all IC (C) and CN units (F).

### Pooling analysis

The finding that IC but not CN responses provided accurate estimates of psychometric threshold and slope (Figure 9A) begs the question of whether and how converging inputs to the IC could provide such psychometric-neurometric correlations. Though previous studies have addressed the question of how IC responses emerge from CN inputs (Nelson and Carney 2004; Hewitt and Meddis 1994), none have used CN data from behaving animals to address how psychometric performance can emerge from population responses. To test this idea, two different kinds of pooling analyses of CN neuronal responses were performed (see Methods): uniform pooling, which indiscriminately samples the population under the assumption that any neuron can contribute to detection performance, and selective pooling based on the lower envelope principle, which states that performance reflects the responses of the most sensitive neurons in the population. Figure 9B shows how simulated populations of CN units exhibit progressively lower median threshold as a function of the number of units in the population. These trends were fit with a power-law function to permit interpolation between different population sizes. Uniform pooling resulted in a median threshold trend that equaled the IC, and behavior, at a population size of about 25 neurons, whereas the selective pooling approach resulted in pooled thresholds approximating behavior using about 10 units. Figure 9C shows how uniform pooling impacted the slopes of neurometric functions of small populations. The median slope of the model increased systematically as the population size increased, suggesting that IC slopes and psychometric slopes could be explained by pooling, however, the variability of the slopes was greater than that observed for the IC. Interpretation of these results and necessary caveats are addressed in the Discussion section.

**Figure 9.**
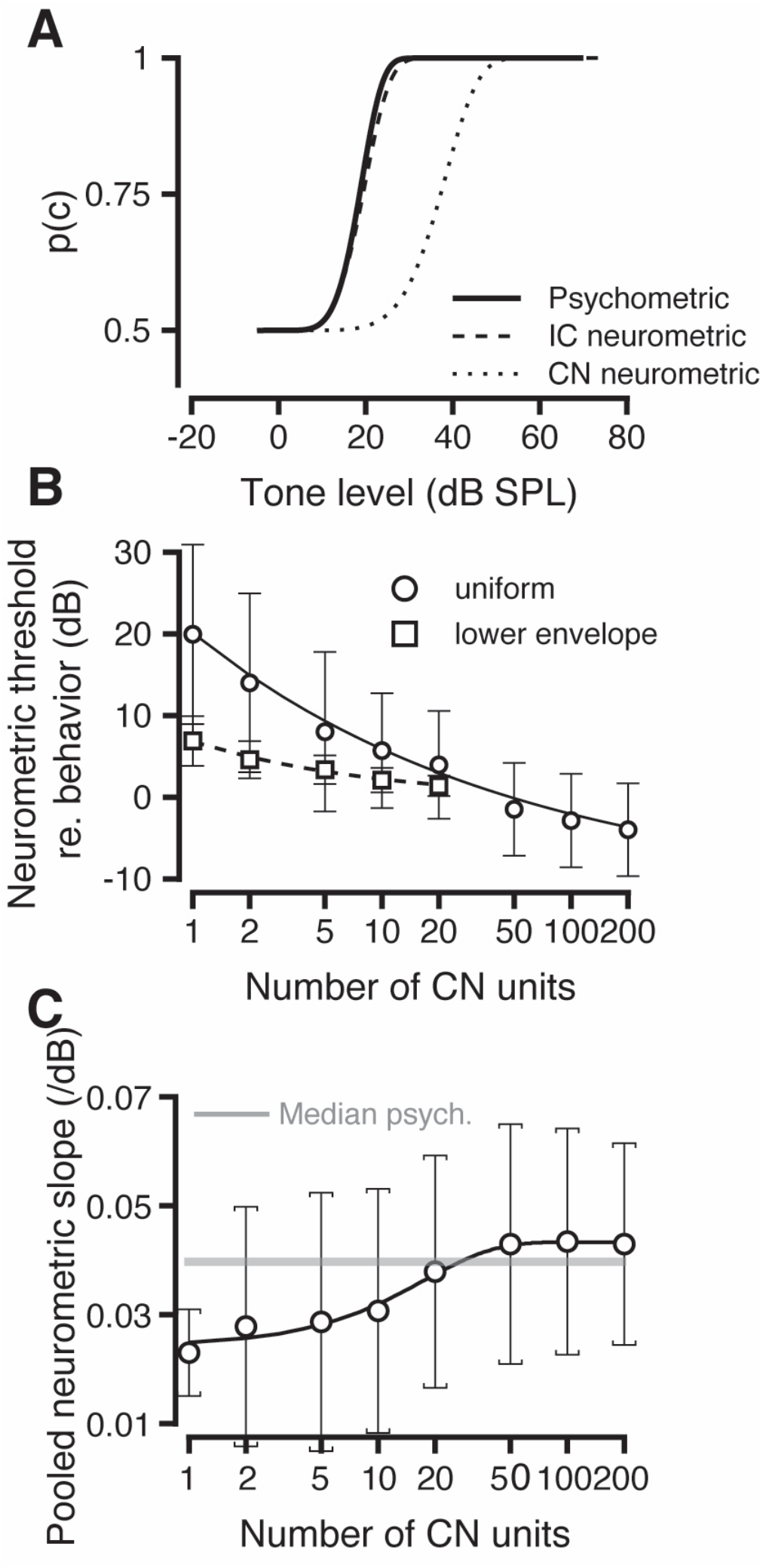
Summary and pooling results. A. Summary of psychometric function (solid lines), IC neurometric functions (dashed), and CN neurometric functions (dotted), illustrating the general finding that IC responses provide better estimates of psychometric threshold and slope than CN responses. B. Plot of the variation of neurometric thresholds predicted by the combination of different numbers of CN units recorded in this study. Uniform pooling (circles) and lower envelope pooling (squares) strategies are shown. Error bars represent 2 SDs. The solid curve and dashed curve represent power-law fits to the uniform (∼ *N*^*-0*.*22*^) and lower envelope pooling (∼ *N*^*-0*.*41*^). C. Plot of the range of neurometric slopes predicted by the uniform pooling of different numbers of CN units. Error bars represent 2 SDs. Median psychometric slope is shown as a grey line.

## Discussion

### Neuronal responses and perception

The results presented here are the first direct neurometric-psychometric correlations reported in the subcortical auditory system. These correlations are similar to studies of cortical regions across sensory modalities (auditory: Lemus et al. (2009b); visual: Britten et al. 1992, 1996; Cook and Maunsell 2002; Purushothaman and Bradley 2005; Palmer et al. 2007; somatosensory: Hernandez et al. 2000; olfactory: Uchida and Mainen 2003), and results in the subcortical vestibular nucleus (Liu et al. 2013). However, they differ in that subcortical neurometric thresholds in the somatosensory system (Vazquez et al. 2012) and visual system (Jiang et al. 2015) did not match psychometric parameters, but IC responses in this study provided excellent estimates of psychometric threshold and slope.

These results also contribute to the understanding of sound detection. The strong match between IC response magnitude and behavioral performance is consistent with other studies showing similarity between behavioral and IC neurophysiological measures (critical bands: Ehret and Merzenich 1985, 1988; Franssen effect: Rajala et al. 2013). However, this does not negate the importance of the cortex for detection (Adams 1986; Heffner and Heffner 1986, 1989, 1990). If the readout of IC information is represented in the cortex, lesions of cortex might disrupt detection. Alternatively, in primates the connections between auditory and motor regions may make the auditory cortex an essential part of the circuit.

Beyond providing insight into readout of auditory information, the present results also relate to theories of information readout derived from other sensory systems, namely that behavior is about as sensitive as the average neuron in visual and somatosensory cortex (Britten et al. 1992; Hernandez et al. 2000). This is in contradiction with the lower envelope principle, which states that the behavioral response is best correlated with the most sensitive neurons (visual cortex: Parker and Newsome 1998; auditory cortex: Johnson et al. 2012). The enhanced sensitivity of the IC relative to the CN begs the question of how neuronal sensitivity in the IC arises from its inputs, which was addressed via pooling (Figure 9B and 9C). The pooling results suggest converging CN inputs can provide accurate estimates of threshold and slope, with pooled CN threshold matching the IC and behavior at a population size of about 10 (lower envelope pooling strategy) or 25 (uniform pooling strategy) units. Anatomically, this pooling is an oversimplification because the IC receives, in addition to CN inputs, inputs from the superior olivary complex, the nuclei of the lateral lemniscus, the contralateral IC, thalamus, and from the cortex (Reviewed in Oliver and Huerta (1992); Schreiner and Winer (2005); and Moore et al. (2010)). However, it provides the first demonstration that small populations in the CN can explain behavior using data from behaving animals and lends support to models that invoke CN-IC convergence to explain IC responses and behavior (Nelson and Carney 2004; Hewitt and Meddis 1994). The finding that small populations of CN neurons can explain behavior is consistent with other work in an amplitude-modulation (AM) discrimination paradigm (Mackey et al. 2022), which is interesting compared to previous pooling studies reporting that similar numbers of primary auditory cortical neurons are required to explain AM detection behavior in macaques (Johnson et al. 2012). This may indicate that perceptual sensitivity attributed to auditory cortex may in fact be inherited from population activity in subcortical regions. The pooling results are presented with the caveat that interneuronal correlations impair the decoding of population activity (Moreno Bote et al. 2014), and due to the recording methods here, interneuronal correlations are not available, an issue that is elaborated upon below.

### Trial-by-trial correlation between neuronal activity and behavioral performance

The covariation between the neuronal response magnitude and behavioral responses on a trial-by-trial basis (Detect probability; Figure 7) are the first reported significant neuronal correlates of behavioral choice in the subcortical auditory system; and one of the few reports of subcortical choice-related activity in any sensory system (Liu et al. 2013). This contrasts previous studies from Kettner and Thompson (1980, 1985), where correlations between neural and behavioral variability were found only outside of the auditory pathway. This difference could be due to species differences (rabbit vs. primate), or more likely, task differences (classical conditioning vs. operant behavior). The DP values reported here for the IC (median =0.647, mean = 0.62) are comparable with the values of choice probability (CP) in the subcortical vestibular nucleus (mean = 0.61, Liu et al. 2013). Interestingly these values from subcortical stations are higher than those reported in studies of primary auditory cortex (Niwa et al. 2012a) and visual cortical area MT during discrimination (mean = 0.555, Britten et al. 1996), and larger than values for primary visual cortex during detection (mean = 0.52, Palmer et al. 2007). One reason that VN neurons have larger CP values could be due to their direct connections with motor nuclei; another possibility that the authors mention is the strong relationship between noise correlation and signal correlation in VN (Liu et al. 2013). While we did not measure noise correlations, one study found no significant noise correlations in the IC of anesthetized guinea pigs in response to speech sounds (Garcia Lazaro et al. 2013). There are currently no descriptions of noise correlation values in CN or IC in awake animals that could compare with this study or highlight the generality of the findings by Liu et al. (2013) for other sensory systems. The emergence of significant choice probabilities in early sensory areas may be due to input from higher order regions (Nienborg and Cumming 2009; Zaidel et al. 2017), and may reflect other non-circuit related factors, such as cognitive state, or the nature of the task (Nienborg et al. 2012). Consistent with this, our group and others have reported task-related modulation of activity in the IC (Rocchi and Ramachandran 2020; Ryan et al. 1984; Shaheen et al. 2021). Another possibility is that response variability in ANF response may introduce a small, statistically non-significant bias between correct and incorrect responses; convergence, such as those in the CN (e.g., Wang and Delgutte 2013), and those between CN and IC, may induce correlated noise that magnifies these effects to reach statistical significance at the IC. Regardless of the underlying circuitry, the evolution of DP over time in the present data suggests it is a rapid process (Figure 7E).

A population of IC units showed significant DP values < 0.5 (their responses on correct trials were lower compared to incorrect trials; Figure 7A). Similar findings have been reported in the visual system: Bosking and Maunsell (2011) report that the DP of some visual neurons (specifically, whose tuning matched the stimulus parameters at the onset of the task) were significantly higher than 0.5; neurons with opposite tuning had DP less than chance. Neurons with DP < 0.5 may perform similar functions as those with DP>0.5. Additionally, the speculation of push-pull action of the populations with DPs above and below 0.5 are intriguing, though the segregating factor was not the monotonicity. Some IC projections to the thalamus are inhibitory (Winer et al. 1996; Peruzzi et al. 1997; Mellott et al. 2014, 2019). The inhibitory and excitatory output neurons could represent the two disparate populations with DP values above and below 0.5. Such convergence could increase the thalamic response contrast between correct and incorrect responses, and may relate to performance monitoring and reward expectation.

The relative paucity of strong correlations between fluctuations in the responses of CN neurons (the first central synaptic station in the auditory system) and behavioral variability is consistent with suggestions of early anatomical work that auditory functions in nonhuman primates are more encephalized (e.g., Adams 1986). The reduced correlations may be only for response magnitude, and not for responses in general; temporal aspects of CN response encoded behavior better than magnitudes (Gai and Carney 2006; Gai et al. 2007). While Gai et al. compared neuronal responses and behavioral responses obtained at different times and in a different species, the possibility of a correlation between behavior and the temporal aspects of CN and IC responses is currently being explored with this and other data sets (Ramachandran, 2018), as well as its implications on circuitry.

These results represent an important step in understanding how subcortical activity relates to auditory perception, how choice-related activity is reflected along the auditory pathway, and provide a baseline for future studies of hearing loss.

## Author Contributions

RR designed the experiments. MD and PB collected data. MD, PB, JG, AH, CAM and JM analyzed the data under RR’s supervision and guidance. RR and CAM wrote and edited the document, the other authors read and edited the document.

## Data availability

Data and code are available upon request. Code is available at https://github.com/seemackey/master

## Acknowledgements

This research was funded by a grant from the National Institutes of Health, RO1 DC 11092. CAM was supported by the Ruth Kirchstein F31 fellowship from NIDCD (F31 DC019823-02). The authors would like to thank Ms. Mary Feurtado and Dr. William Vaughan for help during surgery, Bruce and Roger Williams for hardware, Drs. Emilio Salinas, Jeff Schall, Richard Heitz, and Mark Wallace for helpful comments on an earlier version of the manuscript, Dr. Steve Lisberger and Mr. Scott Ruffner for the use of the FigureComposer software, and Ms. Laura Trice, and Drs. Troy Hackett and Jon Kaas for help with histology.

